# Comprehensive dataset of shotgun metagenomes from stratified freshwater lakes and ponds

**DOI:** 10.1101/2020.11.12.379446

**Authors:** Moritz Buck, Sarahi L. Garcia, Leyden Fernandez Vidal, Gaëtan Martin, Gustavo A. Martinez Rodriguez, Jatta Saarenheimo, Jakob Zopfi, Stefan Bertilsson, Sari Peura

**Author notes:** corresponding authors: Sari Peura, Moritz Buck.

## Abstract

Stratified lakes and ponds featuring steep oxygen gradients are significant net sources of greenhouse gases and hotspots in the carbon cycle. Despite their significant biogeochemical roles, the microbial communities, especially in the oxygen depleted compartments, are poorly known. Here, we present a comprehensive dataset including 267 shotgun metagenomes from 41 stratified lakes and ponds mainly located in the boreal and subarctic regions, but also including one tropical reservoir and one temperate lake. For most lakes and ponds, the data includes a vertical sample set spanning from the oxic surface to the anoxic bottom layer. The majority of the samples were collected during the open water period, but also a total of 29 samples were collected from under the ice. In addition to the metagenomic sequences, the dataset includes environmental variables for the samples, such as oxygen, nutrient and organic carbon concentrations. The dataset is ideal for further exploring the microbial taxonomic and functional diversity in freshwater environments and potential climate change impacts on the functioning of these ecosystems.

## Background & Summary

Stratified lakes are a typical feature of the northern landscapes and are also significant sources of greenhouse gas (GHG) emissions^1^. These lakes largely reside in regions critically impacted by climate change^2^ and the future contribution of these lakes to climate change via GHG emissions is dependent on the microorganisms inhabiting their waters^3,4^. In this regard, organisms residing in the anoxic compartment of these waters are of particular interest, as many of the more potent GHGs are produced under such conditions^1^. However, our knowledge regarding the diversity and functioning of these microbial communities is still sparse, and only a few studies have addressed the ecology and functional features of microorganisms in anoxic lake compartments^5–9^. To address this gap in knowledge, we have collected a set of 267 samples from 41 waterbodies including thermally stratified lakes and ponds from boreal and subarctic regions, as well as a depth profile of a tropical reservoir in Puerto Rico and a time series of depth-resolved samples from a temperate and seasonally stratifying eutrophic lake in Switzerland (Fig. 1, Table 1). For the majority of the lakes, samples were available from across the water column, including the oxic epilimnion, the oxygen transition zone (metalimnion) and the deep anoxic hypolimnion (Supplementary Table S1). Additionally, diverse environmental factors were analysed for all samples, including but not limited to, oxygen, nutrients and organic carbon concentrations (Supplementary Table S1). For all samples, metagenomes were sequenced using deep shotgun sequencing on the Illumina NovaSeq platform at the Science for Life Laboratory (Uppsala University, Uppsala, Sweden). Additionally, for two of the waterbodies (Alinen Mustajärvi and Lomtjärnan), genomes from single cells were amplified and sequenced, specifically targeting poorly known community members, such as members of the lineage *Chlorobia* and candidate phyla radiation. Our goal was to collect a comprehensive dataset that would allow broad analyses of the functioning of the microbial communities in oxygen stratified lakes with emphasis on lakes representing high carbon concentration but with variable environmental conditions with regards to nutrient concentrations, trophic state and other lake features. Based on this data, it is possible to describe the taxonomic identities and genome-encoded functional traits of the predominant microbes across boreal and subarctic lakes and ponds. The dataset represents a major asset for advancing our understanding of the biogeochemical and ecological functioning of these key environments and their present and possible future roles in elemental cycles. The dataset as such can be used, for example, for assessing general metabolic potential of the total lake communities. Further, we have assembled and binned the data into 12665 metagenome-assembled genomes (MAGs), which were further clustered into 3640 metagenomic Operational Taxonomic Units (mOTUs; species level genome clusters). Of these mOTUs, 328 are classified down to species level, while 2141, 878, 220, and 65 could only be assigned to genus, family, order, and class level, respectively. This dataset can, thus, be used for the exploration of population genomes, to study individual community members, and for deeper genome characterization of poorly known members of the resident communities, including their metabolic potentials.

**Table 1.** Location, maximum depth, number of samples and sampling years for all of the lakes included to the dataset.

| site name | country | latitude | longitude | max depth<br>sampled | number<br>of<br>samples | sampling years |
| --- | --- | --- | --- | --- | --- | --- |
| SAS B1_2 | Canada | 55.23 | -77.7 | 0.45 | 3 | 2014; 2017 |
| SAS B4 | Canada | 55.22 | -77.7 | NA | 2 | 2014; 2017 |
| SAS C2_4 | Canada | 55.23 | -77.7 | 0.75 | 3 | 2014; 2017 |
| SAS C5 | Canada | 55.23 | -77.7 | NA | 2 | 2014; 2017 |
| SAS F5 | Canada | 55.23 | -77.69 | 2.35 | 1 | 2014; 2017 |
| SAS G1 | Canada | 55.23 | -77.7 | 0.15 | 2 | 2014; 2017 |
| SAS H1 | Canada | 55.23 | -77.7 | NA | 1 | 2014; 2017 |
| SAS I1_2 | Canada | 55.23 | -77.7 | 0.35 | 2 | 2014; 2017 |
| SAS I3 | Canada | 55.23 | -77.7 | NA | 2 | 2014; 2017 |
| SAS2A | Canada | 55.23 | -77.7 | 2.25 | 4 | 2014; 2017 |
| SAS2B | Canada | 55.23 | -77.7 | 2.05 | 4 | 2014; 2017 |
| SAS2C | Canada | 55.23 | -77.7 | 1.85 | 4 | 2014; 2017 |
| SAS2D | Canada | 55.23 | -77.69 | 2.35 | 4 | 2014; 2017 |
| Alinen Mustajärvi | Finland | 61.21 | 25.11 | 6.56 | 42 | 2014-2015;2018 |
| Keskinen Rajajärvi | Finland | 61.22 | 25.22 | 10 | 5 | 2015 |
| Lovojärvi | Finland | 61.08 | 25.03 | 13.9 | 7 | 2015 |
| Mekkojärvi | Finland | 61.23 | 25.14 | 3.6 | 51 | 2011-2015 |
| Valkea Kotinen | Finland | 61.24 | 25.06 | 4.5 | 4 | 2015 |
| Ylinen Rajajärvi | Finland | 61.22 | 25.21 | 5 | 4 | 2015 |
| La Plata | Puerto Rico | 18.33 | -66.24 | 15 | 8 | 2018 |
| Bengtsgolen | Sweden | 67.92 | 20.37 | 5.5 | 4 | 2018 |
| Björntjärnan | Sweden | 58.7 | 16.19 | 0.2 | 1 | 2018 |
| Erken | Sweden | 64.12 | 18.78 | 8 | 8 | 2018 |
| Fyrsån | Sweden | 59.84 | 18.64 | 20 | 12 | 2018 |
| Glimmingen | Sweden | 59.8 | 18.51 | 0.1 | 1 | 2018 |
| Haukijärvi | Sweden | 57.93 | 15.57 | 0.4 | 1 | 2018 |
| Kiruna 754378-169136 | Sweden | 67.65 | 21.17 | 3.1 | 4 | 2018 |
| Kiruna Ki1 | Sweden | 67.93 | 20.36 | 2.7 | 4 | 2018 |
| Långsjön | Sweden | 59.64 | 16.14 | 0.2 | 1 | 2018 |
| Lillsjön | Sweden | 63.35 | 14.46 | 3.5 | 21 | 2018 |
| Lomtjärnan | Sweden | 59.86 | 17.94 | 0.1 | 1 | 2016-2018 |
| Lotsjön | Sweden | 59.96 | 17.28 | 0.1 | 1 | 2018 |
| Lumpen | Sweden | 60.04 | 17.56 | 0.1 | 1 | 2018 |
| Malstasjön | Sweden | 59.77 | 18.64 | 0.1 | 1 | 2018 |
| Nästjärnen | Sweden | 64.15 | 18.8 | 5 | 4 | 2018 |
| Odrolstjärnen | Sweden | 63.39 | 14.46 | 4.4 | 2 | 2018 |
| Parsen | Sweden | 58.34 | 16.2 | 0.2 | 1 | 2018 |
| Plåten | Sweden | 59.86 | 18.54 | 0.1 | 1 | 2018 |
| Stortoveln | Sweden | 57.93 | 15.55 | 0.2 | 1 | 2018 |
| Loclat | Switzerland | 47.01 | 6.59 | 9 | 44 | 2008-2009 |
| Toolik | USA | 68.63 | -149.6 | NA | 4 | 2017 |

**Table 2.**
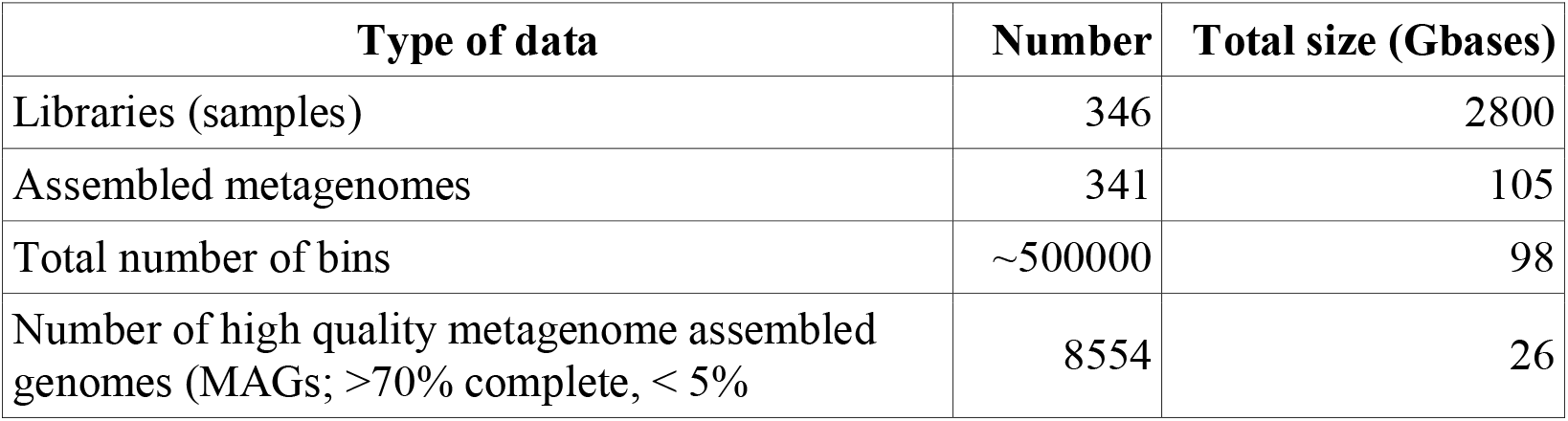
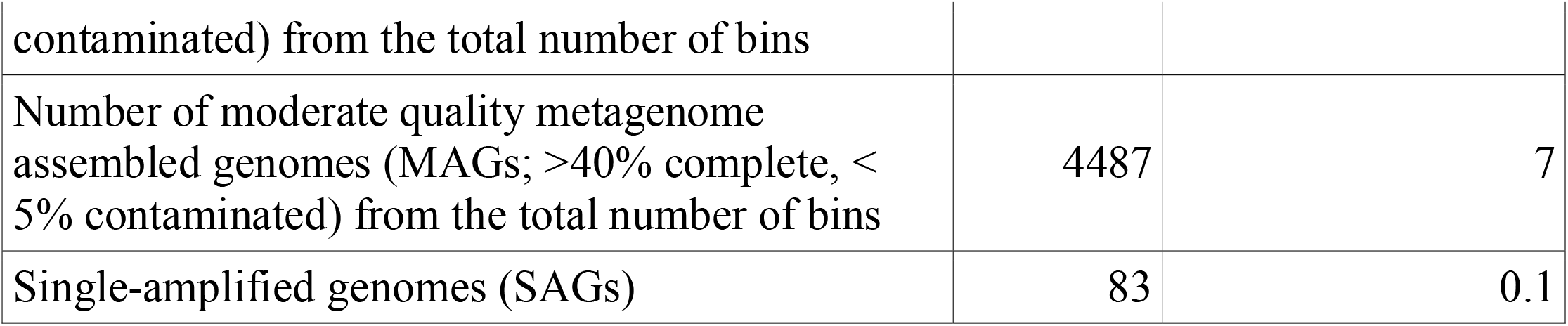
Summary of all genomic data.

**Figure 1.**
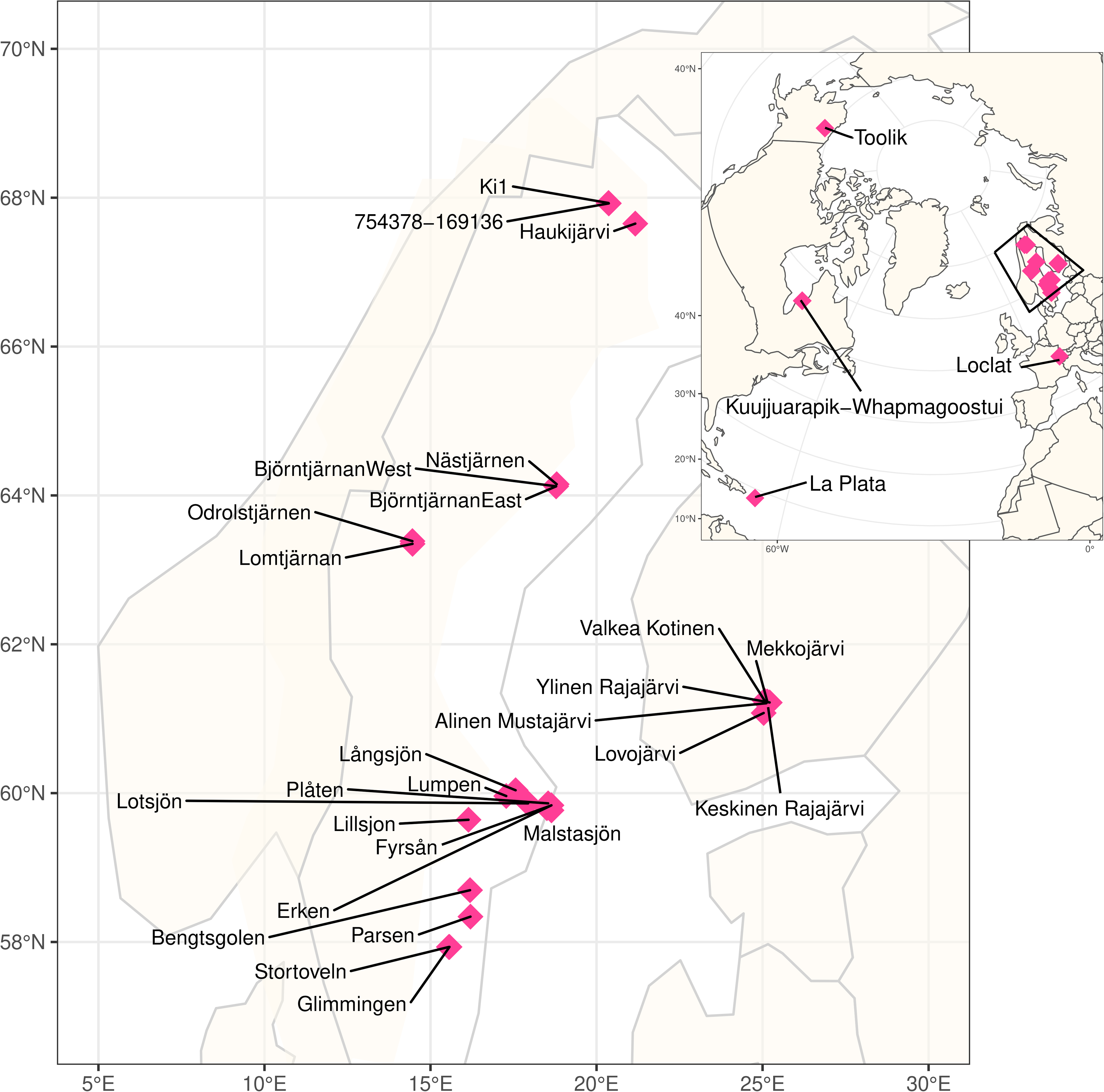
Global distribution of sampling sites. Large Scandinavian map represents the squared region in the insert.

## Methods

### Sample collection

The 267 samples were collected between 2009 and 2018 from 41 locations expanding from the subarctic region to the tropics (Fig. 1, Supplementary Table S1) and processed using the same analytical pipeline (Fig. 2). The majority of the samples were collected using a depth-discrete Limnos tube-sampler (Limnos, Poland), with the exception of the samples from La Plata reservoir (Puerto Rico), which were collected using horizontal Van Dorn sampler (5L capacity) and samples from Lake Loclat, which were collected using a deployed PVC-inlet connected to a peristaltic pump via tubing. Of all the lakes, 29 were sampled during the open water season and the majority of the lakes were sampled once. For 12 of the lakes only surface samples taken during the ice-covered period in winter were available, and one of the Swedish lakes (Lake Lomtjärnan) was sampled twice during the ice-covered period. Moreover, a total of 5 samples (one depth profile) from the time series of the Swiss lake (Loclat) were taken from under the ice. Time series samples were taken for Lake Loclat (seven time points, Supplementary Table S1) and for Lake Mekkojärvi (22 time points, see Saarenheimo et al.^10^ for details). For most lakes and ponds, samples were collected from multiple depths, including samples from the oxic surface layer (epilimnion), the layer with steepest change in oxygen concentration and temperature (metalimnion) and from the layer where oxygen levels were below the detection limit (hypolimnion). The exception to this were the 12 Swedish lakes sampled during ice-covered period, and five shallow ponds in Canada, for which only one sample from the oxic surface layer was taken (see Supplementary Table S1).

**Figure 2.**
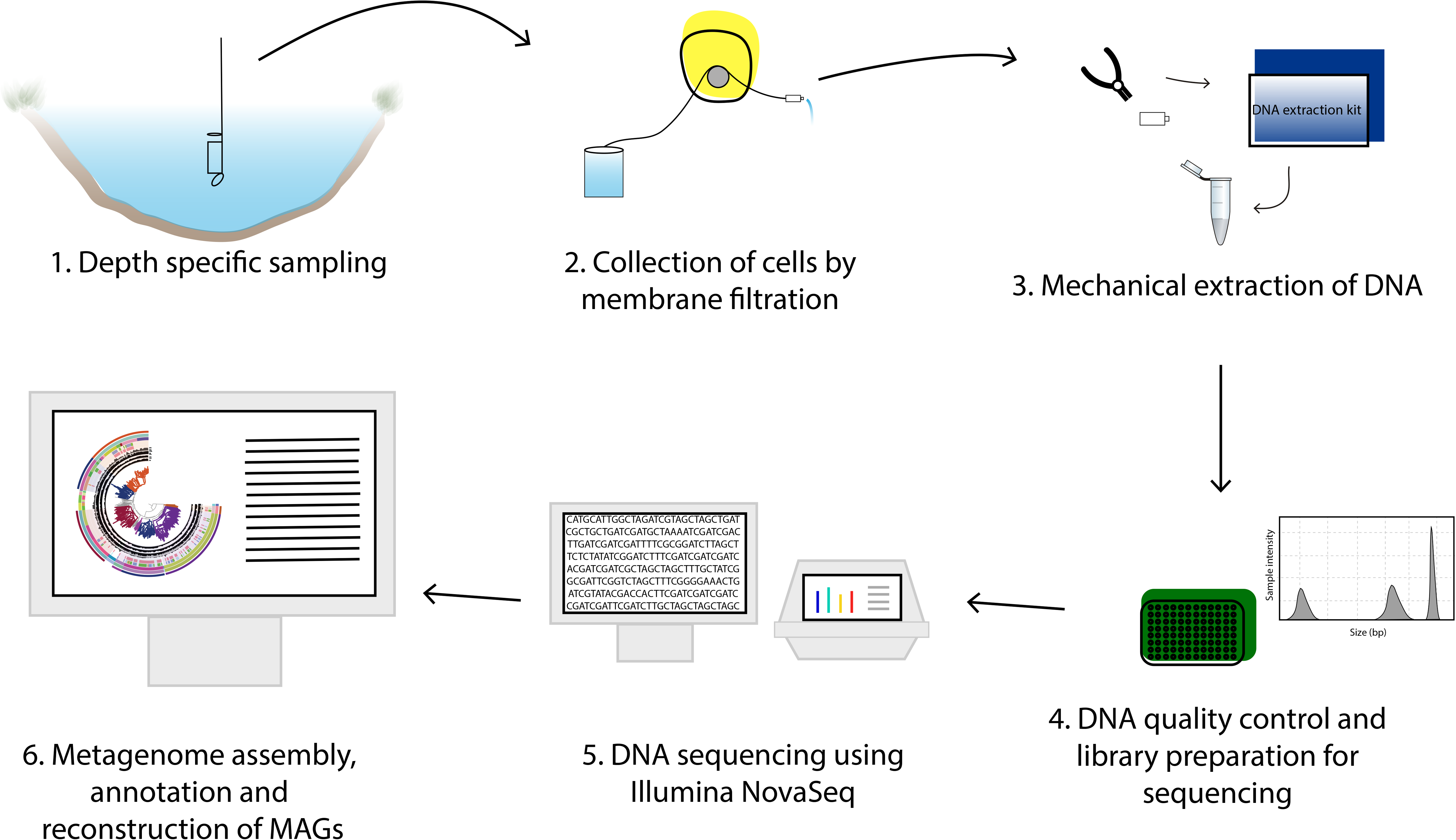
Overview of the workflow from sample collection to mOTUs.

From two of the lakes, Lake Lomtjärnan in Sweden and Lake Alinen Mustajärvi in Finland, samples were collected also for single cell sorting. From both locations samples were preserved in glycerol-TE (gly-TE) and from Lomtjärnan samples were preserved also using phosphate buffered saline (PBS). For both preservants, the samples were flash frozen in liquid nitrogen after first incubating for 1 minute at ambient temperature.

Simultaneous to collection of the DNA samples, also samples for environmental variables were taken. Variables included temperature, pH, conductivity, oxygen, total and dissolved nutrients (P and N species), gases (CO_2_ or dissolved inorganic carbon and methane (CH_4_)), total or dissolved organic carbon, iron, sulfate and chlorophyll a (Supplementary Table S1 and Supplementary Table S2 for the methods). As the samples were collected during multiple years and by different research groups, there was some variation for the procedures between the different sampling occasions, leading to variation in the final set of environmental data across the samples.

### DNA extraction and metagenome sequencing

Most of the DNA samples were collected on 0.2 μm Sterivex filters (Millipore), except for the time-series samples collected from Loclat, which were collected by vacuum filtration onto 47 mm polycarbonate membrane filters with 0.2 μm pore size, and time series samples from Finnish Lake Mekkojärvi, for which the water for DNA extraction was collected from epilimnion (0-0.5 m), metalimnion (0.5-1 m) and hypolimnion (1-3 m) and pooled samples from each stratum were stored in 100 ml plastic containers and frozen at −20°C and eventually freeze-dried (Alpha 1-4 LD plus, Christ). For all filter samples, water was filtered until the filter clogged. All filters were stored frozen (−20 to −80 °C) until the extraction of DNA. For all samples, DNA was extracted using PowerSoil DNA extraction kit (MoBio, Carlsbad, CA, USA) following the manufacturer’s instructions.

Sequencing libraries were prepared from 10 or 20 ng of DNA using the ThruPLEX DNA-seq Prep Kit according to the manufacturer’s preparation guide. Briefly, the DNA was fragmented using a Covaris E220 system, aiming at 400 bp fragments. The ends of the fragments were end-repaired and stem-loop adapters were ligated to the 5’ ends of the fragments. The 3’ end of the stem loop were subsequently extended to close the nick. Finally, the fragments were amplified and unique index sequences were introduced using 7 cycles of PCR followed by purification using AMPure XP beads (Beckman Coulter).

The quality of the libraries was evaluated using the Agilent Fragment Analyzer system and the DNF-910-kit. The adapter-ligated fragments were quantified by qPCR using the Library quantification kit for Illumina (KAPA Biosystems/Roche) on a CFX384Touch instrument (BioRad) prior to cluster generation and sequencing. The sequencing libraries were pooled and subjected to cluster generation and paired-end sequencing with 150 bp read length S2/S4 flow-cells and the NovaSeq6000 system (Illumina Inc.) using the v1 chemistry according to the manufacturer’s protocols.

Base calling was done on the instrument by RTA (v3.3.3, 3.3.5, 3.4.4) and the resulting .bcl files were demultiplexed and converted to fastq format with tools provided by Illumina Inc., allowing for one mismatch in the index sequence. Additional statistics on sequence quality were compiled with an in-house script from the fastq-files, RTA and CASAVA output files. Sequencing was performed by the SNP&SEQ Technology Platform in Uppsala, Sweden.

### Single-cell sorting and DNA amplification

All Gly-TE cryopreserved samples were thawed and diluted in 1 xPBS if needed while all plates with PBS were UV-treated with a dose of 2 J prior to sorting. Samples collected from both lakes were sorted, and then screened for organisms belonging to candidate phyla radiation. Samples collected from Lake Lomtjärnan were additionally subjected to sorting based on autofluorescence to identify and sequence cells belonging to lineage *Chlorobia*.

For obtaining SAGs from representatives of the candidate phyla radiation, samples were first stained with 1 x SYBR Green I for approximate 30 minutes. Subsequent single cell sorting was performed with a MoFlo Astrios EQ (Beckman Coulter, USA) cell sorter using a 488 nm laser for excitation, 70 μm nozzle, sheath pressure of 60 psi and 0.1 μm sterile filtered 1 x PBS as sheath fluid. Individual cells were deposited into empty 384-well plates (Biorad, CA USA) UVed at 2 Joules using a CyCloneTM robotic arm and the most stringent single cell sort settings (single mode, 0.5 drop envelope). Green fluorescence (488-530/40) was used as trigger and sort decisions were made based on combined gates of 488-530/40 Height log vs 488-530/40 Area log and 488-530/40 Height log vs SSC with increasing side scatter divided up in three different regions. Flow sorting data was interpreted and displayed using the associated software Summit v 6.3.1. Next, individual cells were subject to lysis, neutralization and whole genome amplification using MDA based on the protocol and workflow described by Rinke et al.^11^ but with several modifications. Reagent mastermixes were added using the MANTIS liquid dispenser (Formulatrix) and the LV or HV silicone chips. The lysozyme, D2 buffer, stop solution and MDA-mastermix were each dispensed with its own chip. Most MDA-reactions were run using the phi29 from ThermoFisher but a few were run with a more heat-stable phi29, EquiPhi also provided by ThermoFisher. The MDA reaction was carried out in a total volume of μl. Thawed, sorted cells were first pre-treated with 400 nl/well of 12 U/μl of Ready-Lyse™ Lysozyme Solution (R1804M, Lucigen) at room temperature for 15 minutes before adding 400 nl Qiagen lysis buffer D2 followed by incubation at 95°C for 10 seconds and 10 minutes on ice. Reactions were neutralized by adding 400 nl Qiagen Stop solution. Four μl of MDA mix containing 1x reaction buffer, 0.4 mM dNTP, 0.05 mM exonuclease-resistant Hexamers, 10 mM DTT, 1.7 U phi29 DNA polymerase (ThermoFisher Scientific) and 0.5 μM Syto13 was added to a final reaction volume of 5.2 μl. All reagents except SYTO13 were UV decontaminated at 2 Joules in a UV crosslinker. The whole genome amplification was run at 30 °C for 7 or 10 h followed by an inactivation step at 65 °C for 5 min. The reaction was monitored in real time by detection of SYTO13 fluorescence every 15 minutes using a FLUOstar^®^ Omega plate reader (BMG Labtech, Germany) or a qPCR instrument. The EquiPhi protocol was run as previously described for ThermoFisher phi29 with the following exceptions; the EquiPhi polymerase was added in 1U/reaction, reaction buffer included with the polymerase was used and the reaction was carried out at 45 °C. The single amplified genome (SAG) DNA was stored at −20°C until further PCR screening, library preparation and Illumina sequencing. SAGs were screened using the bacterial PCR primers targeting the 16 S rRNA gene, Bact_341 F and Bact_805 R^12^. The reactions were run in a LightCycler 480 PCR machine (ROCHE, MA USA) in 10 μl and a final concentration of 1 x LightCycler480 SYBR Green I Master mix, 0.25 μM of each primer and 2 μl of 60 to 80 times diluted SAGs. Following a 3 min denaturation at 95°C, targets were amplified for 40 cycles of 95 °C for 10 s, 55 °C for 20 s, 72 °C for 30 s and a final 10 min extension at 72 °C followed by melting curve analysis. The products were purified using the NucleoSpin Gel and PCR clean-up purification kit (Macherey-Nagel, Germany), quantified using the Quant-iT TM PicoGreen^®^ dsDNA assay kit (Invitrogen, MA USA) in a FLUOstar^®^ Omega microplate reader (BMG Labtech, Germany) and submitted for identification by Sanger sequencing at Eurofin Genomics. All SAGs were further screened using the newly designed primers targeting the phylum Parcubacteria 684F-OD1 (3’ GTAGKRRTRAAATSCGTT 5’) and 784R (5’ TAMNVGGGTATCTAATCC -3’). These primers target with good specificity 67% of Parcubacteria in the SILVA database^13^. Parcu-PCR was run at 3 min at 95 °C, 40 cycles of 95 °C for 10 s, 55 °C for 20 s, 72 °C for 30 s and a final 10 min extension at 72 °C followed by melting curve analysis. The products were purified using the NucleoSpin Gel and PCR clean-up purification kit (Macherey-Nagel, Germany), quantified using the Quant-iT TM PicoGreen^®^ dsDNA assay kit (Invitrogen, MA USA) in a FLUOstar^®^ Omega microplate reader (BMG Labtech, Germany) and submitted for identification by Sanger sequencing at Eurofin Genomics.

To recover *Chlorobia* single amplified genomes, sorting was done in 2016 on a MoFlo™ Astrios EQ sorter (Beckman Coulter, USA) using a 488 and 532 nm laser for excitation, a 70 μm nozzle, a sheath pressure of 60 psi, and 0.1 μm filtered 1x PBS as sheath fluid. An ND filter ND=1 and the masks M1 and M2 were used. The trigger channel was set to the forward scatter (FSC) at a threshold of 0.025% and sort regions were defined on autofluorescence using laser 532 nm and band pass filters 710/45 and 664/22. Three populations were sorted based on differences in autofluorescence signals. The sort mode was set to single cell with a drop envelope of 0.5. The target populations were sorted at approximately 400 events per second into 96-well plates containing 1 μl 1x PBS per well with either 1 or 10 cells (positive control) deposited. A few wells remained empty (no cell sorted) were kept as negative controls. Sorted plates were stored frozen at −80 °C.

The subsequent whole genome amplification was performed in 2018 using the REPLI-g Single Cell kit (QIAGEN) following the instructions provided by the manufacturer but with total reaction volume reduced to 12.5 μl. The denaturation reagent D2, stop solution, water, and reagent tubes and strips were UV-treated at 2.5 J. The lysis was changed slightly to 10 min at 65 °C, followed by 5 min on ice before adding the stop solution. To the master mix containing water, reaction buffer, and the DNA the polymerase we added SYTO 13 (Invitrogen) at a final concentration of 0.5 μM. The amplification was performed at 30 °C for 8 hours in a plate reader with fluorescence readings every 15 min. The reaction was stopped by incubating it for 5 min at 65 °C. The plate was stored for less than a week at −20 °C. Amplified DNA was mixed thoroughly by pipetting up and down 20 times before diluting it 50x and 100x in nuclease-free water. The DNA was screened for bacterial 16S rRNA applying the primers Bact_341 F (5’- CCTACGGGNGGCWGCAG- 3’) and Bact_805 R (5’-GACTACHVGGGTATCTAATCC-3’)^12^ using the LightCycler^®^ 480 SYBR Green I Master (Roche) kit. The PCR mix contained 1.5 μl diluted amplified DNA, 1x the LightCycler^®^ 480 SYBR Green I Master mix, 0.25 μM of each primer, and nuclease-free water in a total reaction volume of 10 μl. The PCR cycling (5 min at 95°C, followed by 40 cycles of 10 sec at 95 °C, 20 sec at 60 °C, 30 sec at 72 °C) was followed by meltcurve analysis on the LightCycler^®^ 480 Instrument (Roche). DNA of confirmed *Chlorobia* was sent to sequencing as outlined below.

### Library Preparation and Illumina sequencing of the single cells

For the candidate phyla radiation-targeted analysis, Illumina libraries were prepared from sixty SAGs mainly selected from the screening procedure in a PCR-free workflow using the sparQ DNA Frag & Library Prep Kit (Quantabio) and IDT for Illumina TruSeq UD Indexes (Illumina). Libraries were prepared from 50-250 ng of MDA-products in 25% of the recommended reaction volumes according to manufacturer’s instructions. The MDA-products were fragmented for 7 minutes (5 minutes for 4 samples) without using the DNA Frag Enhancer Solution. Library insert sizes were determined using Bioanalyzer High Sensitivity DNA Kit (Agilent). Each library was quantified using the KAPA Library Quantification kit (Roche) in 5 □l reaction volumes in a 384-well plate run on LightCycler 480 (Roche) to allow equimolar pooling before sequencing on Illumina HiSeqX v2.5 PE 2 × 150 bp. For the *Chlorobia*-targeted sequencing, amplified DNA from 23 SAGs were quantified individually with Qubit dsDNA HS assay kit (ThermoFisher Scientific) and diluted to 0.2 ng/ul in nuclease free water. Sequencing libraries were prepared with Nextera XT DNA Library Preparation Kit and combinatorial combinations of molecular identifiers in the Nextera XT Index Kit (Illumina, CA USA) according to manufacturer’s instructions. Libraries with an average length of 1200 bp were quantified with Qubit dsDNA HS assay kit to allow pooling of equal amounts of the libraries based on mass. The libraries were sequenced on an Illumina MiSeq v3 PE 2 x 300 bp.

### Data processing of the metagenome and single cell sequences

The metagenome sequencing resulted in a total of ~10^7^ paired-end reads of length 2×150 bp, amounting to a total of total 3 Tbp. The raw data was trimmed using Trimmomatic (version 0.36; parameters: ILLUMINACLIP:TruSeq3-PE.fa:2:30:10 LEADING:3 TRAILING:3 SLIDINGWINDOW:4:15 MINLEN:36)^14^ (Supplementary Table S3). The trimmed data was assembled using Megahit (version 1.1.13)^15^ with default settings. Two types of assemblies were done, single sample assemblies for all the samples individually and a total of 53, mainly lake-wise, co-assemblies (see Supplementary Table S4), some samples of the Canadian ponds have also been coassembled with previously sequenced libraries of the same sample (see Supplemental Table S5). The relevant quality controlled reads were mapped to all the assemblies using BBmap^16^ with default settings and the mapping results were used to bin the contigs using Metabat (version 2.12.1, parameters --maxP 93 --minS 50 -m 1500 -s 10000)^17^. Genes of obtained bins were predicted and annotated using Prokka (version 1.13.3)^18^ using standard parameters except for the bin containing all the unbinned contigs where the –metagenome flag was used. Single-cell libraries were processed similarly to the metagenomes, but without the binning step, and using the single-cell variant of the SPAdes^19^ assembler instead of Megahit.

Prokaryotic completeness and redundancy of all bins from Metabat and for all assembled single cells were computed using CheckM (version 1.0.13)^20^ (Supplementary Table S6 and S7 for MAGs and SAGs, respectively). Average Nucleotide Identity (ANI) for all bin-pairs was computed with fastANI (version 1.3)^21^. The bins were clustered into metagenomic Operational Taxonomic Units (mOTUs) starting with 40% complete genomes with less than 5% contamination. Genome pairs with ANI above 95% were clustered into connected components. Additionally, less complete genomes were recruited to the mOTU if its ANI similarity was above 95%. Bins were taxonomically annotated in a two-step process. GTDB-Tk (version 102 with database release 89)^22^ was used first with default settings. Using this classification an lca database for SourMASH (version 1.0)^23^ was made. This database as well as one based on the GTDB release 89 was then used with SourMASH’s lca classifier for a second round of classification of bins that were not annotated with GTDB-tk (Supplementary Table S8).

The taxonomic diversity of the bacterial (Fig. 3) and archaeal (Fig. 4) mOTUs, respectively, were visualized in a tree format. The trees were computed using GTDB-tk with one representative MAG per mOTU of the stratfreshDB, and one random representative genome per family of the GTDB. Trees were visualized using anvi’o^24^.

**Figure 3.**
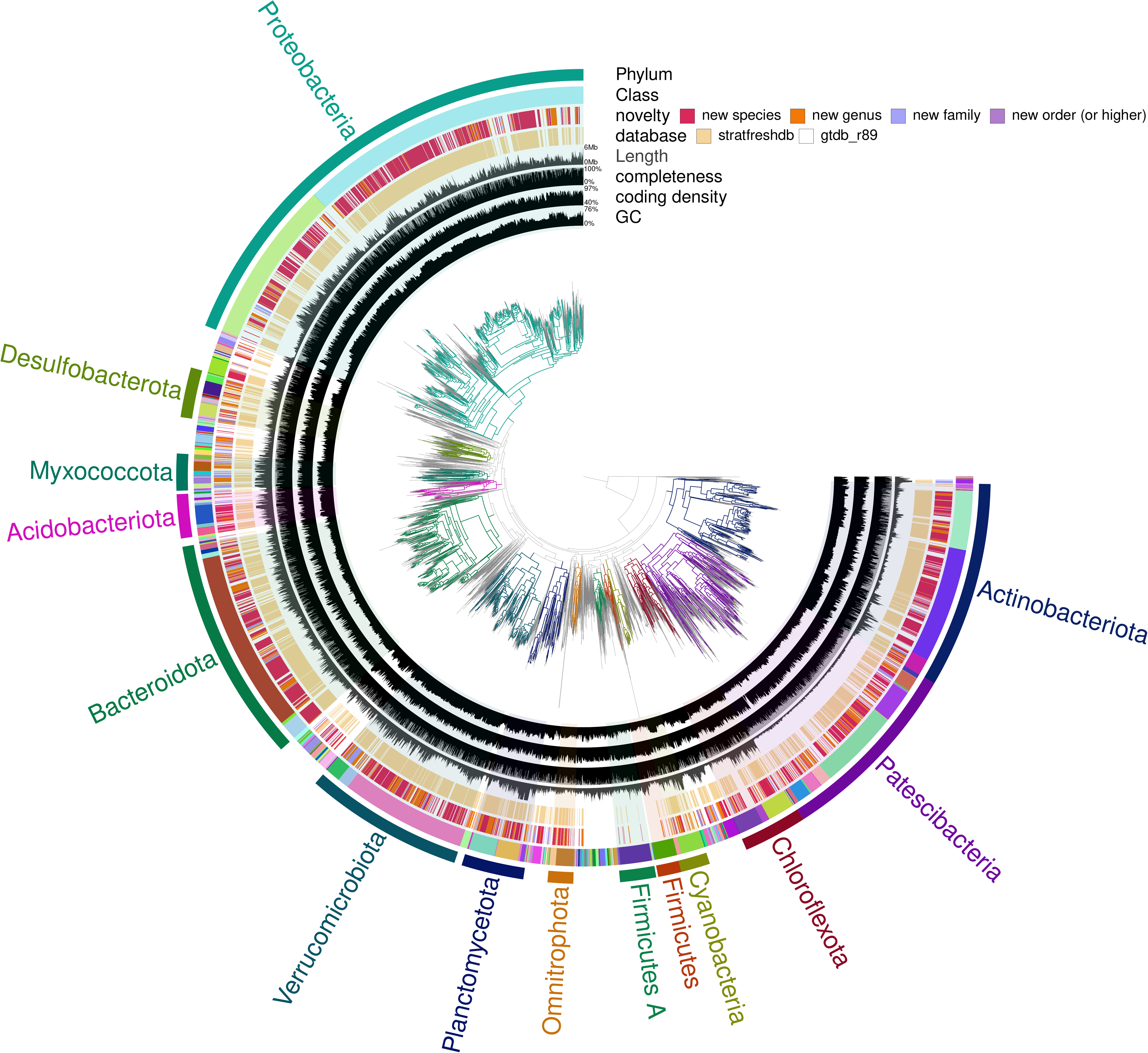
Bacterial diversity of the stratfreshDB. Interactive version with more information available at https://anvi-server.org/moritzbuck/bacterial_diversity_of_the_stratfreshdb.

**Figure 4.**
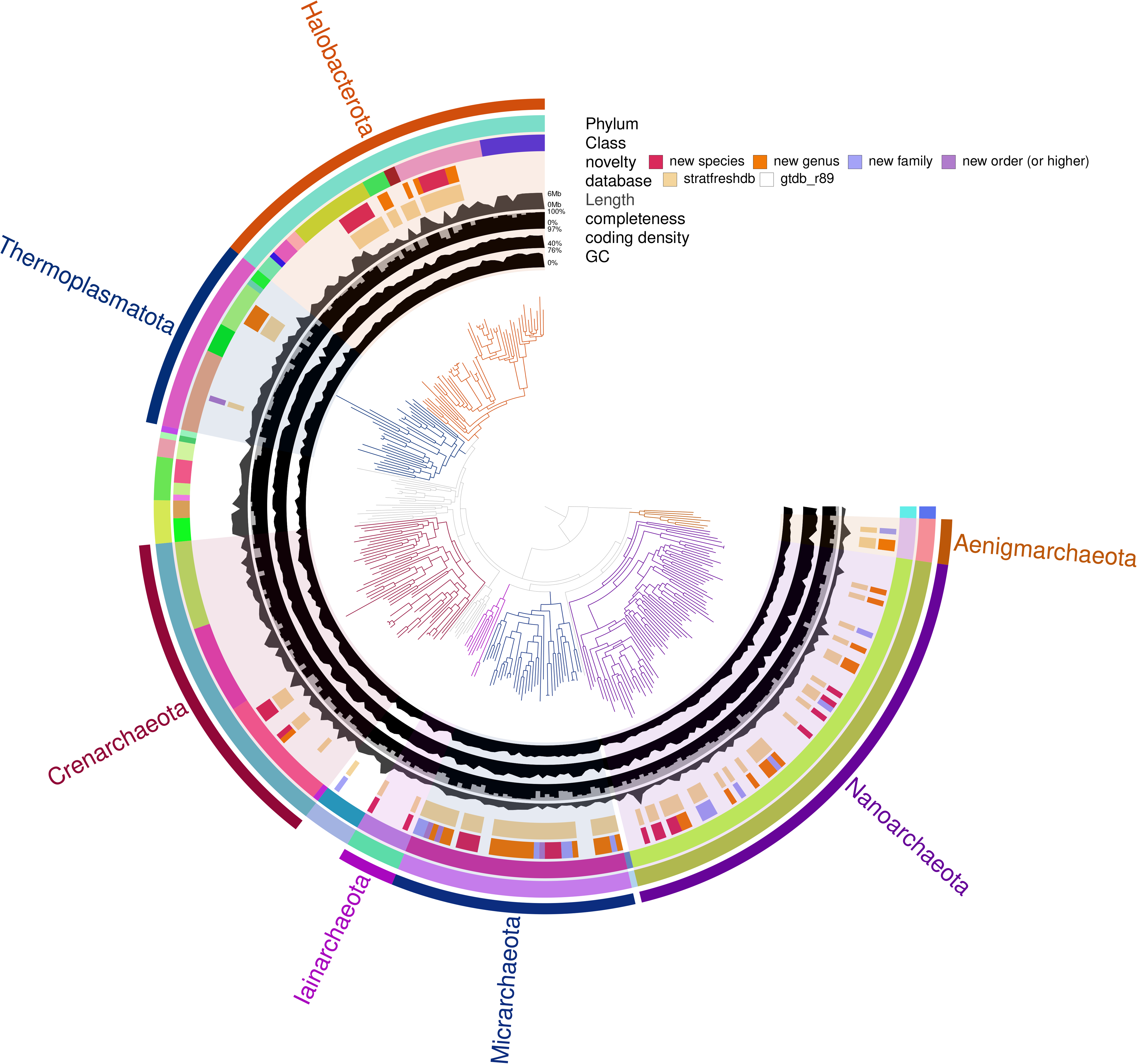
Archaeal diversity of the stratfreshDB. Interactive version with more information available at https://anvi-server.org/moritzbuck/archaeal_diversity_of_the_stratfreshdb.

## Data Records

All sequences are deposited to the European Nucleotide Archive (ENA, mirrored to SRA, and accessible at the NCBI) under the project number PRJEB38681. Statistics for reads, metagenome assemblies, high-quality bins (e.g. MAGs), and SAGs can be found in Supplementary Table S3, S2, S4, and S5 respectively. Additional tables and information can be found under the doi: 10.17044/scilifelab.13005311

## Technical Validation

Reads were QCed by the sequencing facility using the MultiQC [doi:10.1093/bioinformatics/btw354] wrapper for FastQC^25^. MAG and SAG completeness and contamination rate have been computed with CheckM^20^.

## Usage Notes

All data submitted to the ENA can also be easily accessed at https://export.uppmax.uu.se/uppstore2018116/stratfreshdb/ using the paths in the files available at doi:10.17044/scilifelab.13005311 for navigation

## Code Availability

Code used to process the data is available at https://github.com/moritzbuck/metasssnake, code for the computing of mOTUs is available at https://github.com/moritzbuck/mOTUlizer and some additional scripts, particularly the script used for the submission of the data is available at https://github.com/moritzbuck/0023_anoxicencyclo.

## Acknowledgements

Milla Rautio and Maxime Wauthy are acknowledged for their help with access and sampling at the CEN research station in Whapmagoostui-Kuujjuarapik (Quebec, Canada). We thank Sergio Morales for his help obtaining the samples from Puerto Rico. We thank Maliheh Mehrshad for moral support. We are grateful to Karólína Einarsdóttir, Hannu Nykänen, Christophe Paul, Carla Perez, Antti Rissanen, Henrique Sawakuchi, Vicente Sedano, Christian Wurzbacher, and numerous other researchers and students for encouragement and help with the sampling. Lammi Biological Station is acknowledged for their help with sampling of the Finnish lakes. The sequencing was funded by a grant from the Science for Life Laboratory biodiversity program and SciLifeLab fellows program. Sequencing was performed by the SNP&SEQ Technology Platform in Uppsala. The facility is part of the National Genomics Infrastructure (NGI) Sweden and Science for Life Laboratory. The SNP&SEQ Platform is also supported by the Swedish Research Council and the Knut and Alice Wallenberg Foundation. The computations were performed on resources provided by SNIC through Uppsala Multidisciplinary Center for Advanced Computational Science (UPPMAX) under Project SNIC snic2020-5-19. We also want to thank the SciLifeLab Data Center for helping us to make the data available more freely. Samplings were funded by misc. grants from the Swedish Research Council, the Swedish Research Council Formas, The K*A Wallenberg foundation, INTERACT, Academy of Finland, and Olsson-Borgh foundation.

## Author contributions

SP, SB, MB and SLG conceived the project idea and secured funding for realizing the sequencing. All authors contributed to the collection of the samples. MB conducted and coordinated the bioinformatic processing of the metagenomic data. LFV, GM, GMR, JS, and JZ have provided considerable number of samples and data. SP drafted the manuscript with significant inputs and contributions from SB, MB and SLG, and assistance of all the other authors. All authors have read and approved the final version of the manuscript.

## Manuscript metadata tables

Supplementary Tables:

Supplementary Table S1. Contextual data of biosamples: Table containing chemical, geographical, and other contextual data about the physical samples collected for this dataset. Contains the fields:

- sample_id: name of the sample, corresponds to sample_name in other tables
- Run: R[123], the data was sequenced in 3 Illumina NovaSeq runs, this is the ID of the run
- site name: name of the location samples (e.g. lake-name, or ID)
- Lake_code: short ID for that location
- region: geographical region the location is in (), or closes large city
- sample_accession: SRA/ENA sample accession
- assembly_accession: ENA metagenome assembly accession
- sample owner: the person responsible for field sampling
- all other columns correspond to metadata-fields of ENA mostly from ENA-checklist ERC000024, the second row of the table is units corresponding to columns

Supplementary Table S2. Methods used for analyses of the environmental data.

Supplementary Table S3. Data about sequenced libraries: table containing statistics, accession number and other contextual data about the actual sequenced libraries, including metagenomic as well as single-cell amplified genome libraries. Contains the fields:

- library_identifier: ENA identifier of library
- total_spots: number of reads
- total_bases: number of nucleotides
- total_size: size of the files in bytes
- title: unique title of library at SRA/ENA
- sample_id: ENA/SRA sample ID
- sample_name: name of the sample
- fwd: relative path to the “.gz”-compressed forward reads, in FASTQ-format, relative to https://export.uppmax.uu.se/uppstore2018116/stratfreshdb/
- rev: relative path to the “.gz”-compressed reverse reads, in FASTQ-format, relative to https://export.uppmax.uu.se/uppstore2018116/stratfreshdb/
- library_type: metagenome if it is a metagenome library, SAG if it is the library of a single-cell amplified genome

Supplementary Table S4. Data about metagenomic assemblies: A table containing statistics, accession number and other contextual data about the 342 metagenomic assemblies computed for this data-set. Contains the fields:

- assembly: name of the assembly, if it is a single-sample assembly, this name is the sample name, else an appropriate name
- length: length of the assembly in bases
- GC: GC fraction of the assembly
- nb_contigs: number of contigs in the assembly
- path: relative path to the “.bz2”-compressed assembly, in FASTA-format relative to https://export.uppmax.uu.se/uppstore2018116/stratfreshdb/
- type: single_sample_assembly or coassembly_assembly, if single_sample_assembly only reads for a single sample have been used to assemble this assembly, else reads from multiple biological samples were used to coassemble this coassembly
- library: ENA/SRA biosample-identifier of read-libraries used, matches the identifiers in Supp. Table S1

Supplementary Table S5. A table containing accession number of a number libraries that had been sequenced previously and have been re-sequenced for this project, in some assemblies, both libraries were used to improved assembly quality

Supplementary Table S6. A table containing statistics, accession numbers and other contextual data about the 12682 Metagenome Assembled Genomes of moderate and high-quality submitted to ENA (e.g. completeness estimated higher then 40% and redundancy lower then 5%). A table with the whole ~500.000 bins generated in this study can be found at the scilifelab-figshare doi:10.17044/scilifelab.13005311.

Contains the fields:

- bin_id: the unique identifier of a MAG
- GC: GC-content of MAG
- coding_density: coding-density estimate, e.g. sum of the length of all predicted amino-acid sequences times 3 divided by the length of the genome (in bases)
- completeness: estimated completeness computed by CheckM
- contamination: estimated redundancy computed by CheckM
- strain_heterogeneity: computed by CheckM
- length: length in bases of the MAG
- nb_contigs: number of contigs in the MAG
- nb_proteins: number of predicted proteins encoded in the bin (for the predicted amino-acid sequences in the easy-access repository)
- gtdbtk_classification: classification by GTDBtk r89
- sourmash_classification: classification by sourmash using 2 databases, the GTDB (r89), and a database based on our GTDBtk classification
- taxonomy: conclusion of the two previous
- mOTU: membership to a metagenomic Operational Taxonomic Unit (mOTU), e.g. species level clustering of bins
- path: relative path a tar-ball containing nucleotide, CDS- and amino-acid sequences, GFF, eggNOGmapper, and sourmash-signature files. Relative to https://export.uppmax.uu.se/uppstore2018116/stratfreshdb/
- accession: SRA/ENA sample ID of the assembly
- genbank_accession: genbank accession number

Supplementary Table S7. A table containing statistics, accession numbers and other contextual data about the Single cell Assembled Genomes submitted to ENA. Contains the same fields as table S6.

Supplementary Table S8. A table containing statistics and other contextual data about the 3640 metagenomic Operational Taxonomic Units (mOTUs), e.g. species-level MAG clusters. Contains the fields:

- mOTU_name: ID of the mOTU
- mean_ANI: mean of all Average Nucleotide Identity of all the MAGs in the cluster
- consensus_tax: predicted consensus taxonomy of the cluster
- mean_good_ANI: mean of all ANIs of all the high-quality MAGs (70%+ completeness, <5% redundancy) in the cluster
- mean_decent_ANI: mean of all ANIs of all the moderate quality MAGs (40%+ completeness, <5% redundancy) in the cluster
- nb_genomes: number of bins in the cluster
- nb_good_genomes: number of high quality MAGs (70%+ completeness, <5% contamination) in the cluster
- nb_decent_genomes: number of moderate quality MAGs (40%+ completeness, <5% contamination) in the cluster
- representative_MAGs: a bin representing the cluster well (very high completeness, very low redundancy)

## References

1 Bastviken, D., Cole, J., Pace, M. & Tranvik, L. Methane emissions from lakes: Dependence of lake characteristics, two regional assessments, and a global estimate. Glob. Biogeochem. Cycle 18, 12, doi:10.1029/2004gb002238 (2004).

2 Blenckner, T. et al. in The impact of climate change on European lakes (ed DG George) 339–358 (Springer Science+Business Media, 2010).

3 Yvon-Durocher, G. et al. Methane fluxes show consistent temperature dependence across microbial to ecosystem scales. Nature 507, 488–491, doi:10.1038/nature13164 (2014).

4 Cavicchioli, R. et al. Scientists’ warning to humanity: microorganisms and climate change. Nature 17, 569–586, doi:10.1038/s41579-019-0222-5 (2019).

5 Peura, S. et al. Novel Autotrophic Organisms Contribute Significantly to the Internal Carbon Cycling Potential of a Boreal Lake. mBio 9, e00916–00918, doi:doi.org/10.1128/mBio.00916-18 (2018).

6 Garcia, S. L., Szekely, A. J., Bergvall, C., Schattenhofer, M. & Peura, S. Decreased Snow Cover Stimulates Under-Ice Primary Producers but Impairs Methanotrophic Capacity. mSphere 4, e00626–00618, doi:10.1128/mSphere.00626-18 (2019).

7 Taipale, S., Jones, R. I. & Tiirola, M. Vertical diversity of bacteria in an oxygen-stratified humic lake, evaluated using DNA and phospholipid analyses. Aquatic Microbial Ecology 55, 1–16, doi:10.3354/ame01277 (2009).

8 He, S., Lau, M., Linz, A., Roden, E. & McMahon, K. Extracellular electron transfer may be an overlooked contribution to pelagic respiration in humic-rich freshwater lakes. mSphere 4, e00436–00418, doi:10.1128/mSphere.00436-18 (2019).

9 Garcia, S., Salka, I., Grossart, H. & Warnecke, F. Depth-discrete profiles of bacterial communities reveal pronounced spatio-temporal dynamics related to lake stratification. Environmental Microbiology Reports 5, 549–555, doi:10.1111/1758-2229.12044 (2013).

10 Saarenheimo, J. et al. Bacterial community response to changes in a tri-trophic cascade during a whole-lake fish manipulation. Ecology 97, 684–693 (2016).

11 Rinke, C. et al. Obtaining genomes from uncultivated environmental microorganisms using FACS–based single-cell genomics. Nature Protocols 9, 1038–1048 (2014).

12 Herlemann, D. P. R. et al. Transitions in bacterial communities along the 2000 km salinity gradient of the Baltic Sea. Isme Journal 5, 1571–1579, doi:10.1038/ismej.2011.41 (2011).

13 Quast, C. et al. The SILVA ribosomal RNA gene database project: improved data processing and web-based tools. Nucleic Acids Res. 41, D590–D596 (2013).

14 Bolger, A., Lohse, M. & Usadel, B. Trimmomatic: A flexible trimmer for Illumina Sequence Data. Bioinformatics 30, 2114–2120, doi:10.1093/bioinformatics/btu170 (2014).

15 Li, D., Liu, C. M., Luo, R., Sadakane, K. & Lam, T. W. MEGAHIT: an ultra-fast single-node solution for large and complex metagenomics assembly via succinct de Bruijn graph. Bioinformatics 31, 1674–1676, doi:10.1093/bioinformatics/btv033 (2015).

16 BBMap short read aligner (University of California, Berkeley, California, 2016).

17 Kang, D. W. D., Froula, J., Egan, R. & Wang, Z. MetaBAT, an efficient tool for accurately reconstructing single genomes from complex microbial communities. Peerj 3, 15, doi:10.7717/peerj.1165 (2015).

18 Seemann, T. Prokka: rapid prokaryotic genome annotation. Bioinformatics 30, 2068–2069, doi:10.1093/bioinformatics/btu153 (2014).

19 Bankevich, A. et al. SPAdes: A New Genome Assembly Algorithm and Its Applications to Single-Cell Sequencing. J. Comput. Biol. 19, 455–477 (2012).

20 Parks, D. H., Imelfort, M., Skennerton, C. T., Hugenholtz, P. & Tyson, G. W. CheckM: assessing the quality of microbial genomes recovered from isolates, single cells, and metagenomes. Genome Res. 25, 1043–1055, doi:10.1101/gr.186072.114 (2015).

21 Jain, C., Rodriguez-R, L. M., Phillippy, A. M., Konstantinidis, K. T. & Aluru, S. High throughput ANI analysis of 90K prokaryotic genomes reveals clear species boundaries. Nature Communications 9, 5114, doi:10.1038/s41467-018-07641-9 (2018).

22 Parks, D. et al. A standardized bacterial taxonomy based on genome phylogeny substantially revises the tree of life. Nature Biotechnology 36, 996–1004, doi:10.1038/nbt.4229 (2018).

23 Brown, C. T. & Irber, L. sourmash: a library for MinHash sketching of DNA. Journal of Open Source Software 1, 27, doi:10.21105/joss.00027 (2016).

24 Eren, A. M. et al. Anvi’o: an advanced analysis and visualization platform for ‘omics data. Peerj 3, e1319 (2015).

25 Andrews, S. FastQC: a quality control tool for high throughput sequence data, <http://www.bioinformatics.babraham.ac.uk/projects/fastqc> (2010).

